# Explainable machine learning-assisted exploration of chromatin dynamics reveals chromosome-specific response to serum starvation

**DOI:** 10.1101/2025.08.08.669316

**Authors:** Taras Redchuk, Antti P. Pennanen, Harri Jäälinoja, Olli Natri, Lassi Paavolainen, Maria K. Vartiainen

## Abstract

Chromatin is dynamic at all length scales, influencing chromatin-based processes, such as gene expression. Even large-scale reorganization of whole chromosome territories has been reported upon specific signals, but lack of suitable methods has prevented analysis of the underlying dynamic processes. Here we have used CRISPR-Sirius for time-lapse imaging of chromatin loci dynamics during serum starvation. We show that chromosome 1 loci move towards the nuclear envelope during the first hour of serum starvation in a chromosome-specific manner. Machine learning-assisted exploration of acquired multiparametric data combined with the Shapley values-based explanation approach allowed us to uncover the critical features that characterize chromatin dynamics during serum starvation. This analysis reveals that although serum starvation affects overall nuclear morphology and chromatin dynamics, chromosome 1 loci display a specific response that is characterized by maintenance of dynamics in constrained environment, and long “jumps” at the nuclear periphery. Interestingly, the two homologous chromosomes display differential behaviors, with the more peripheral homolog being more responsive to the signal than the internal one. Overall, the presented machine learning-assisted dataset exploration helps us navigate the multidimensional data to understand the underlying dynamic processes and can be applied to a wide variety of research questions in imaging and cell biology in general.

## INTRODUCTION

The human genome is non-randomly organized within the cell nucleus. This complex, multilevel organization from wrapping the DNA around histones as nucleosomes to chromatin loops and topologically associated domains (TADs) and finally to chromosome territories ensures genome functionality at all stages of the cell cycle and in response to intrinsic and extrinsic cues. In addition to folding, also the localization within the 3D nucleus influences chromatin functions. Radial organization, the relative proximity to the generally repressive nuclear envelope, is one measure that can be used to describe the nuclear localization of specific chromatin loci and has been shown in many examples to correlate with not only gene activity, but also for example replication timing. In addition to the nuclear envelope, also proximity to nuclear bodies, such as nucleolus and nuclear speckles, can influence chromatin organization and functions (Bizhanova and Kaufman, 2020).

An essential feature of chromatin organization, at all organizational levels, is its dynamics. The polymer nature and the different interactions of chromatin result in subdiffusive behavior that is more restricted than Brownian motion (Dion and Gasser, 2013). However, observations of specific loci switching to what could be described as active motion were reported as early as the late 90s (Bornfleth *et al*., 1999). Subsequent transgene-based approaches revealed long-range repositioning of specific chromosome loci in response to transcriptional activation (Chuang *et al*., 2006; Dundr *et al*., 2007; Khanna *et al*., 2014). Interestingly, interfering with actin and myosin activity prevented repositioning of the loci. Actin machinery has also been linked to chromatin dynamics especially during DNA damage response by moving damaged DNA to favorable repair sites (Caridi *et al*., 2018; Schrank *et al*., 2018). Thus, while chromatin generally displays subdiffusive dynamics, also long-range, actin-dependent movement has been described.

The particular position of each chromosome remains relatively stable during interphase, forming what is known as a chromosome territory (Walter *et al*., 2003). Nevertheless, studies using mainly fluorescent in situ hybridization (FISH) techniques have revealed large-scale repositioning of chromosome territories in response to serum removal (Mehta *et al*., 2010), DNA damage (Mehta *et al*., 2013; Kulashreshtha *et al*., 2016) and senescence (Mehta *et al*., 2021), as well as during various disease and differentiation contexts (Williams *et al*., 2002; Matarazzo *et al*., 2007). In primary human fibroblasts, serum withdrawal results in chromosome repositioning of about half of the chromosomes in a process that requires energy and the activity of an actomyosin motor. The repositioning is rather rapid, taking place in about 15 minutes, as well as reversible (Mehta *et al*., 2010). Since these studies have been performed in fixed cells, the underlying dynamic processes have remained unclear.

Although both FISH- and transgene-based approaches have advanced chromatin dynamics studies, they still retain several key disadvantages. FISH-based techniques do not allow following the cells live, thus losing the data on real-time motility. In turn, the transgene-based systems require the introduction of an engineered construct into the genome. This restricts the ability to observe the dynamics in untouched cells in a non-invasive way and leaves no freedom to choose the endogenous locus for study. CRISPR-based approaches utilizing endonuclease deficient Cas9 (Chen *et al*., 2013; Ma *et al*., 2016) effectively address both issues, providing precise non-invasive localization of a sequence-of-choice in live cells. Indeed, for example CRISPR-Sirius system, where an engineered guide RNA (gRNA) scaffold amplifies the signal for live imaging (Ma *et al*., 2018) has revealed cell-cycle dependent chromatin dynamics (Ma *et al*., 2019). Live imaging of chromatin dynamics results in multidimensional data that has the potential to reveal mechanistic insights into the underlying biological process. Nevertheless, many previous studies concentrate on a limited set of dynamic properties, such as mean squared displacement (MSD)-related metrics (Dion and Gasser, 2013), necessitating the development of data analysis tools to fully capitalize on the information-rich imaging data.

In this study, we combined CRISPR-Sirius for sequence-specific live visualization of chromatin with machine learning-assisted dataset exploration. Explanation of the trained classifiers’ decisions using a game theory-rooted approach, named SHAP (Lundberg and Lee, 2017), allowed us to extract the patterns underlying chromosome 1 loci dynamics upon serum starvation. This manifests as fast spontaneous jumps rather than continuous movement in a particular direction. In parallel, serum starvation induces a global change in morphological characteristics of the nucleus, such as decrease in the area, which, however, does not explain the radial reorganization of chromosome 1 loci. Finally, under starvation, homologous chromosomes exhibit differential behavior, with loci on peripheral homologs often moving faster than central ones. We are confident that, together with the emerging advanced techniques tackling the complexity of particle tracking (Pinholt *et al*., 2021) and other multimodal biological data (Sturma *et al*., 2023), explainable ML-assisted dataset exploration will add to a toolbox allowing an information-efficient and unbiased data analysis applicable far beyond the showcased scenario.

## RESULTS

### Monitoring chromosome relocalization upon serum starvation with CRISPR-Sirius

Relocalization of chromosomes has previously been described upon DNA damage, heat shock and serum starvation (Abdel-Halim *et al*., 2006; Mehta *et al*., 2010; Mehta *et al*., 2013; Kulashreshtha *et al*., 2016). These studies have been carried out in fixed cells, precluding analysis of the dynamic processes involved. To tackle this issue, we have used CRISPR-Sirius, a tool enabling live imaging of specific chromatin loci (Ma *et al*., 2018), to study chromosome relocalization upon serum starvation. To establish the feasibility of this approach, we first performed a prescreening in fixed U2OS cells that were grown in serum containing media (10%) or serum starved (0.3%) for 1h. We focused on the set of chromosomes previously suggested to display differential radial position between proliferating and quiescent nuclei of human dermal fibroblasts or upon serum starvation (Mehta *et al*., 2010), namely chromosomes 1, 10, and 13, as well as X as a control (Supplementary Figure S1A). Of the imaged chromosomes, chromosome 1 loci displayed detectable relocalization towards the nuclear periphery, manifested also as change in the skewness of distance to edge distribution (Fig. 1A,B). Relocalization was less evident for the rest of the screened chromosomes. To assess the relocalization of chromosome 1 similarly as in the conventional FISH-based assay (Mehta *et al*., 2010; Mehta *et al*., 2021), we also analyzed its nuclear localization by dividing the nucleus into concentric shells with an equal area. Also this approach confirmed the relocalization of chromosome 1 loci towards the nuclear periphery (Fig. 1C). The inability to detect changes in radial positioning of the other chromosomes’ loci may be due to the differential labeling process or the used cell line. Chromosome paints used in the FISH assays cover large parts of the chromosome, whereas CRISPR-Sirius targets only specific repetitive loci within the chromosome. Moreover, here we use the osteosarcoma cell line U2OS, whereas the original studies employed human dermal fibroblasts, and it is possible that different cell lines display distinct chromatin-based responses to serum starvation. Nevertheless, the pre-screening tests showed that CRISPR-Sirius-based tools can be used to study relocalization of chromosome 1 upon serum starvation, opening the possibility for dynamic studies in living cells.

**Figure 1:**
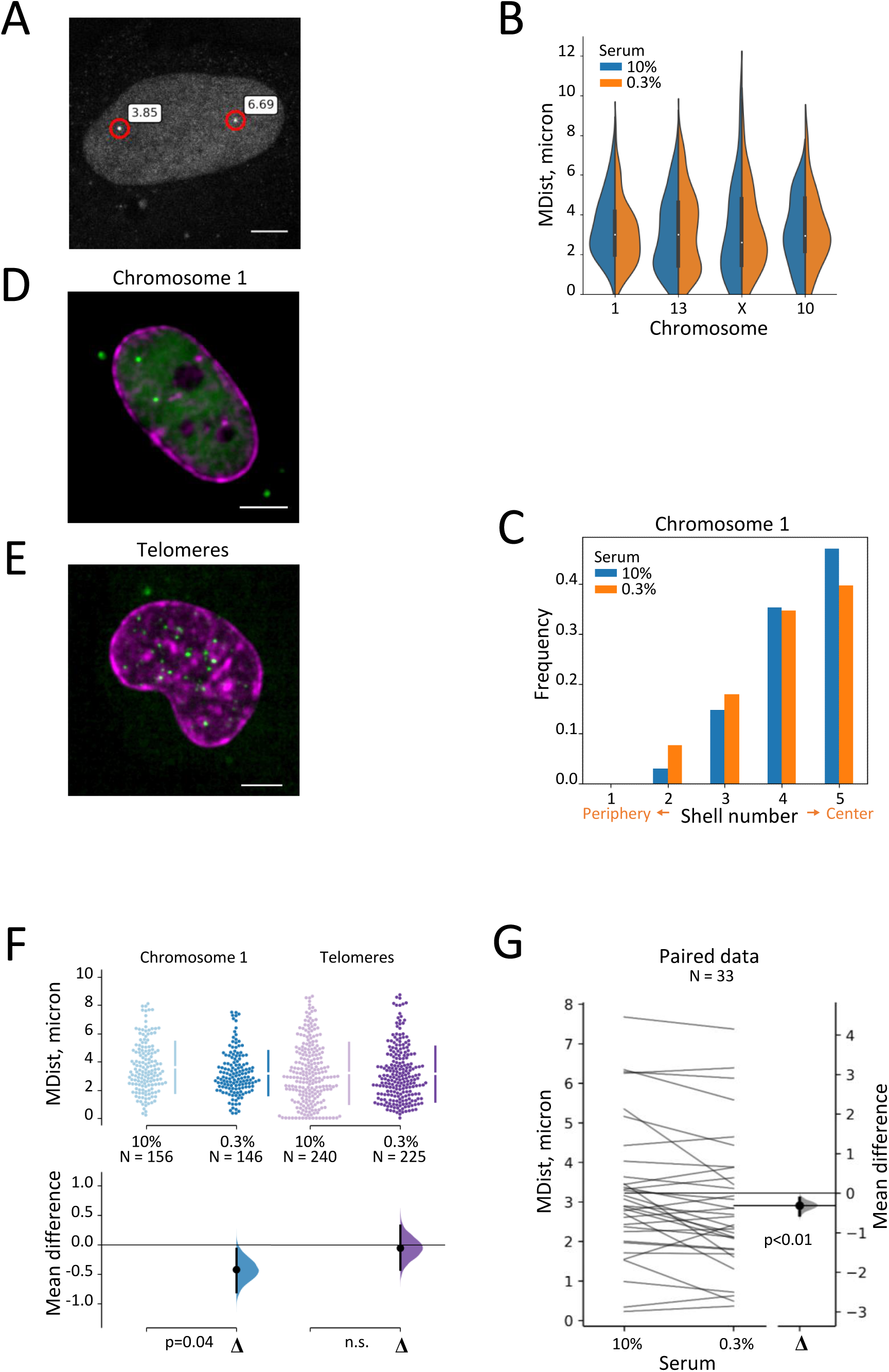
Chromatin dynamics upon serum starvation with CRISPR-Sirius. **A**, Representative image of a nucleus in fixed sample with chromosome loci visualized using CRISPR-Sirius. Minimal distance to edge indicated in white boxes, microns. Scale bar 6 microns. **B**, Kernel density estimate distributions showing the minimal distance to the nuclear edge (MDist) for different chromosome loci in serum (10%) or after 1h serum starvation (0.3%). **C**, Fractions of chromosome 1 loci detected in shells of equal volume derived from the division of the nuclear area by sequential morphological erosion. **D,E** Representative images of U2OS cells expressing CRISPR-Sirius plasmids with the gRNA targeting chromosome 1 (**D**) and telomeres (**E**) in green, and LaminA-mCherry for nucleus segmentation in magenta. Scale bar as above. **F,** Minimal distance to edge (MDist) of visualized loci in serum (10%) and under starvation (0.3%). Each dot in a swarmplot, upper panel, represents individual locus; number (N) of analyzed loci indicated below the swarmplot. Lower panel represents the distributions of bootstrapped delta mean values for datasets. Error bars show 95% confidence intervals in both upper and lower panels. Statistical significance with Mann-Whitney U rank test. **G**, Chromosome 1 loci are observed to localize closer to the nuclear periphery under starvation, also when measured in the same cells before and after (within 40 min) medium change. Every line represents the change in minimal distance to the nuclear periphery (MDist) for a particular locus; left scale. The distribution shows bootstrapped delta mean values for paired dataset; right scale. Error bars show 95% confidence intervals. The difference is significant, paired two-tailed t-test, normality Shapiro-Wilk (α=0.05), p<0.01.

### Live imaging confirms locus-specificity of chromosome 1 radial relocalization upon starvation

To analyze the dynamic behavior of chromosome 1 loci in response to serum starvation, the cells were transfected with CRISPR-Sirius and LaminA-mCherry for nuclear segmentation (Fig. 1D). Randomly chosen cells were imaged in 10% FBS and in low serum (0.3% FBS), within the first 40 min into starvation. The minimum distance to the edge (MDist; all measured parameters and corresponding naming are summarized in Table 1) was calculated for each frame. As a control, we monitored the movement of telomeres (Fig. 1E) in order to distinguish locus-specific effects of serum starvation from general nucleus-level rearrangements. For chromosome 1 loci, ∼ 0.5 micron shift towards nuclear periphery was observed in live cells under serum starvation compared to cells grown in serum (Fig. 1F, Supplementary Table 1 for all statistics). While telomeres were generally more variable in terms of their radial distribution, their averaged minimal distance to the nuclear edge did not show any significant change under starvation, with the differences of the means being close to zero (Fig. 1F). To exclude possible sampling bias before and after medium change, we performed a set of experiments with the same cells imaged before and after changing the media to low serum (Fig. 1G, Supplementary Figure S1B,D). Here, a statistically significant shift in mean distance (*MDist*, Table 1) of 0.32 microns towards the periphery was observed in starvation (Fig. 1G). When examining only the first twenty minutes after medium change, the effect was even more pronounced in this paired dataset (Supplementary Figure S1C).

**Table 1:** Features extracted from the imaging data collected in this study.

| **Raw features describing motility, morphology and circular organization** |  |  |
| --- | --- | --- |
| Locus displacement | D | Distance in microns between two consecutively detected locus positions. Represents the speed of the locus movement. |
| Nuclear area | A | Nuclear area in square microns. Together with perimeter describes nuclear size and morphology. |
| Nuclear perimeter | P | Nuclear perimeter in arbitrary units. Shows nuclear shape, higher for irregular nuclei and lower for round ones. |
| Locus distance to nuclear periphery | Dist | Minimal distance from locus to nuclear rim in microns; describes the radial localization of the locus. |
| Change of direction | dTheta | The smaller angle between two consecutive displacements, shows the change in direction of locus movement. Describes directionality of the movement, which is often associated with active transport. |
| **Prior knowledge - driven feature engineering** |  |  |
| Mean locus displacement | MD | Mean of time series, calculated for each locus. Represents the speed of the locus movement. |
| Maximal locus displacement | MaxD | Maximal displacement within time series. Longest step on the track. |
| Variance of locus displacement | VarD | Variance of locus displacement within time series. Heterogeneity in step lengths of the locus trajectory, showing if movement has consistent speed (low VarD, constrained diffusion) or varies widely with occasional jumps (high VarD). |
| Mean nuclear area | MA | Nuclear area, mean of time series, calculated for each nucleus. |
| Mean nuclear perimeter | MP | Nuclear perimeter, mean of time series, calculated for each nucleus. |
| Mean locus distance to nuclear periphery | MDist | Locus distance to periphery, mean of time series, calculated for each locus, see Dist (above). |
| Variance of locus distance to nuclear periphery | VarDist | Variance of locus distance to nuclear rim within time series. Shows if association with the nuclear periphery is stable or varies within the timelapse. |
| Relative variance of locus displacement | rVarD | Variance of locus displacement divided by mean displacement ($\text{VarD} / \text{MD}$). Interpreted as VarD (above), but in relation to general locus speed. |
| Relative variance of locus distance to nuclear periphery | rVarDist | Variance of locus distance to nuclear periphery divided by Mean distance to periphery ($\text{VarDist} / \text{MDist}$). Interpreted as VarDist, but in relation to general radial position of the locus. |
| Total locus displacement | TD | Distance from first to last coordinate of track. Shows how far the locus traveled during the imaging. |
| Persistence of locus movement | Pers | Total locus displacement divided by track length. Shows how directional the movement is. Directionality is usually associated with active transport. |
| Fastest locus (True or False) | ifFast | True for the fastest (highest MD) homologous locus in the nucleus, and False for the others. Used to study, if the homologs behave differently. |
| Range of locus distance to nuclear periphery | DistR | Difference between maximal and minimal value of locus distance to nuclear periphery. Change in radial position within the nucleus; sign indicates the direction (towards or away from periphery). |
| Total locus radial displacement | TDist | Change in locus distance to nuclear periphery, last position minus first. Same as DistR, but absolute. |
| Central locus (True or False) | ifCentr | True for the homologous locus furthest from nuclear rim (highest Dist) and False for the others. Used to study, if the homologs behave differently. |
| Trend of change of locus distance to nuclear periphery | sDist | The slope of linear approximation of locus distance to nuclear periphery change. Shows, if the locus drifts closer to, or further away from the nuclear periphery. |
| Trend of change of nuclear area | sA | The slope of linear approximation of nuclear area change. Shows, if the nucleus is growing or shrinking. |
| Trend of change of nuclear perimeter | sP | The slope of linear approximation of nuclear perimeter change. Describes the dynamics or nuclear shape together with sA. |
| Mean displacement + 2SD outliers (count) | out2sd | Number of displacements 2*SD longer than MD. Shows the number of long jumps on the track. |
| Mean displacement + 3SD outliers (count) | out3sd | Number of displacements 3*SD longer than MD. Same as out2sd, but with a higher threshold for jump length. |

To scrutiny the observed positional changes, we performed an additional control experiment that is conceptually similar to previously utilized for single molecule tracking (Banaz *et al*., 2019) For this, we imaged a series of fixed samples of U2OS cells expressing CRISPR-Sirius plasmids, using an optical setup identical to the one used for live cell imaging. Next, we processed the acquired timelapse images through our localization detection pipeline. Since the cells were chemically fixed, the localization changes obtained represent static error, with the uncertainty originating from optical component vibrations, camera sensor noise, sample photobleaching, non-deterministic analysis steps, etc. Median displacement length detected for fixed sample control was below 20 nm, while changes reported in live imaging peaked at around 80 nm (Supplementary Figure S1E). On the same scale, the change in minimal distance to the nuclear edge, reported in our study under serum starvation in live samples (0.32 and 0.5 micron), is more than one order of magnitude above the static error (Supplementary Figure S1E). This shows the sufficient spatial resolution of the chosen imaging modality.

Thus, chromosome 1 loci relocalize towards the nuclear periphery upon changing the media to contain low serum. This reorganization is specific to chromosome 1 loci and is not detectable when analyzing telomeres.

### Machine learning augmented analysis of timelapse-derived data

To fully capitalize on this methodology that allows for the real-time monitoring of loci position and nuclear morphology in live cells, we have further enhanced the acquired dataset by incorporating both manually engineered features derived from tracking, as well as the raw morphology data (Table 1). Deciphering the mechanisms underlying a particular biological process includes the choice of descriptive features of the process in question, extracting the patterns in features and their interactions and, finally, interpretation of these patterns. In biological imaging these steps usually imply data pre-processing, semantic segmentation, tracking, and modeling (Cohen, 2017). Naturally, each of these steps leads to the generation of new higher-level features. This feature space expansion, boosted by addition of complex features, derived from two or more naïve system characteristics, leads to inadequacy of human interpretation due to combinatorial explosion (Arganda Carreras and Andrey, 2017; Berisha *et al*., 2021) and multiple comparisons problem (Curran-Everett, 2000). To mitigate this problem while maximizing the information gain, we built our analysis pipeline around machine learning-assisted dataset exploration (Fig. 2A). For this, we designed a set of machine learning models and trained them on features extracted from chromosome 1 live imaging dataset (Table 1). A gradient boosting classifier (GBC) and a multilayer perceptron (MLP), relatively simple models working robustly on tabular datasets of this size, were trained on all the features aggregated over time, including the engineered ones. Alternatively, InceptionTime, a deep-learning model architecture designed for time-series classification, was chosen to analyze the raw time-series, without an access to prior knowledge-based engineered features (Fig. 2A).

**Figure 2:**
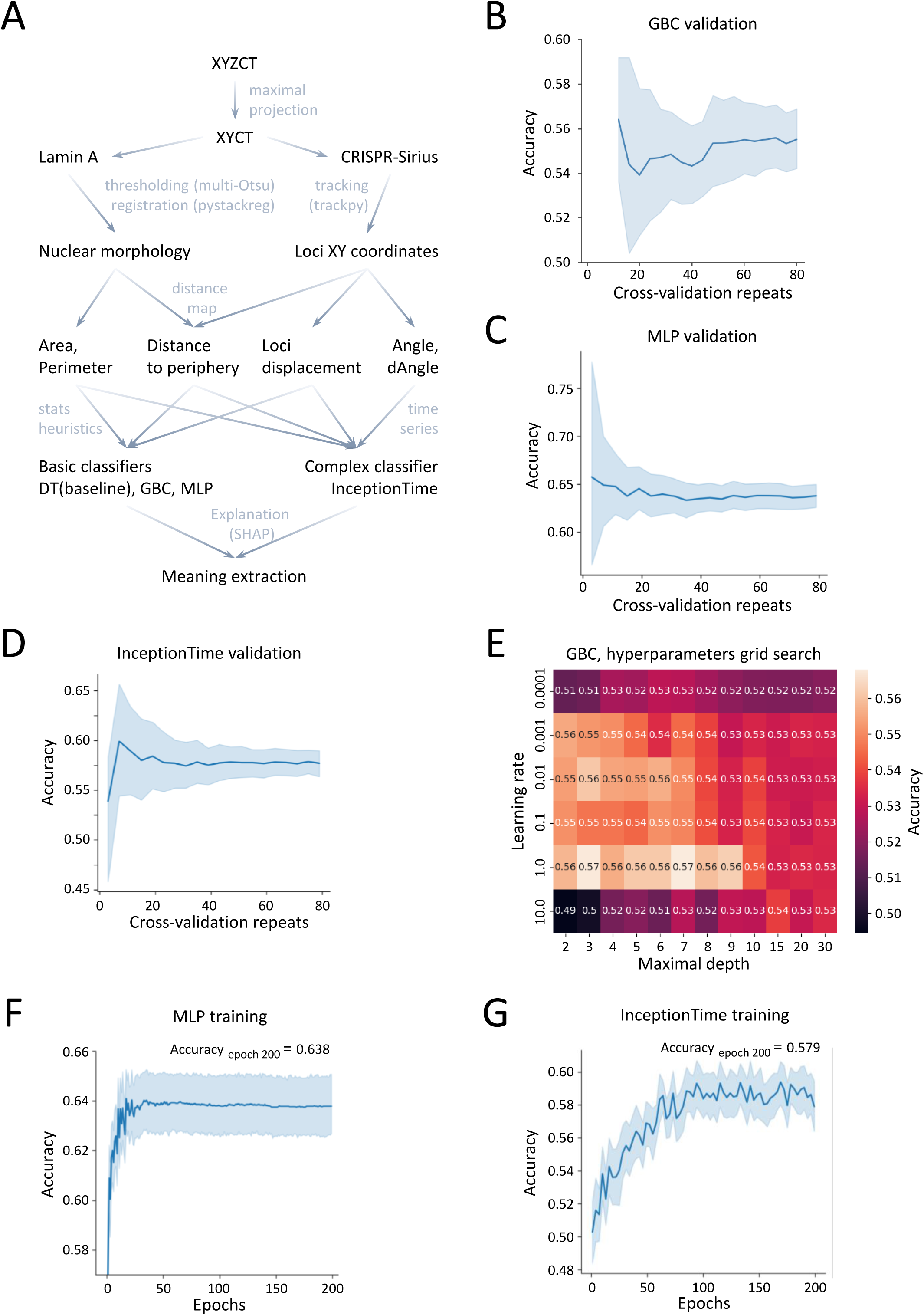
Machine learning-assisted dataset exploration. **A**, Information flow from raw pixel intensities to inference and explanations. (**B-D**), Decrease of accuracy estimation variance in repeated cross-validation of gradient boosting classifier (GBC) (**B**), multilayer perceptron (MLP) (**C**) and InceptionTime (**D**). Shaded area shows 95% confidence interval. **E**, Hyperparameter tuning for GBC. **F**, The accuracy of multilayer perceptron converges after approximately 50 epochs of training at validation accuracy of 0.64; **G**, InceptionTime demonstrated performance comparable to GBC, but without access to manually engineered features. (**F,G**) Shaded area shows 95% confidence interval.

In order to collect the maximum of information from the available finite body of data while being able to assess the classifiers performance, we performed repeated cross-validation of trained models. To validate each model, 20 iterations of random 4-fold splitting were performed, with the loci from the same nucleus always allocated to the same fold to prevent data leakage due to shared morphology features. This allowed a considerable decrease in confidence intervals in sets of estimated model accuracies (Fig. 2B-D).

Next, having only minor class imbalance (Zero Rate = 0.52), we used one level decision tree that basically thresholds one feature at a time, to estimate baseline performance, which reached an accuracy of 0.54 using the same repeated cross-validation strategy as the rest of classifiers. Remarkably, gradient boosting classifier and multilayer perceptron showed higher mean accuracy (0.57 and 0.64; Fig. 2E,F), indicating the ability of the more complex models to gain additional information by abstracting from the feature interactions and non-linearity. After training on the same dataset, InceptionTime demonstrated accuracy of 0.58 (Fig. 2G), exceeding the baseline performance at a level comparable to GBC. Noteworthy, this level was achieved without an access to manually engineered features.

### Model decision explanation by SHAP

In order to extract biological insights from the dataset, the rules used by the machine learning model for inference can be extracted in a process called explanation. The basic way to draw the conclusions about the dataset from model performance would be the assessment of feature importances. However, these techniques often return only absolute importance values, do not allow local explanation, and do not specify the interaction component of the features (Saarela and Jauhiainen, 2021). In contrast, a game theory-based approach named SHAP (SHapley Additive exPlanations (Lundberg and Lee, 2017)) returns the locally relevant additive feature impact values, showing whether the feature contributes to or against a particular class and to what extent.

We used SHAP to explain the classifiers’ decisions and to assess feature interactions for exploratory purposes (Fig. 3A-D). Several features with continuous, non-discrete values, such as mean nuclear area (*MA*) and trend of its change (*sA*), demonstrated high importance and evident correlation between the feature value and its SHAP value in all trained classifiers (Fig. 3A,B,F). Interestingly, correlation between speed (see *D* and *MD*) and its impact on model decision was clear in InceptionTime explanation only; the same is true for distance to edge (*Dist* and *MDist*) (Fig. 3F). The features with discrete values describing outliers from displacement length distribution (*out3sd* and *out2sd*), also showed clear value/SHAP value correlation in GBC and, more prominently, in MLP (Fig. 3A,B), while not used for InceptionTime training. Binary features subclassing homologous chromosomes based on speed or radial position (*ifFast* and *ifCentr*), did not affect GBC, but demonstrated an impact on MLP decisions that is comparable to other features including morphological (Fig. 3B). In accordance with this, discrete and binary features grew in importance in MLP explanation compared to GBC (Fig. 3E).

**Figure 3:**
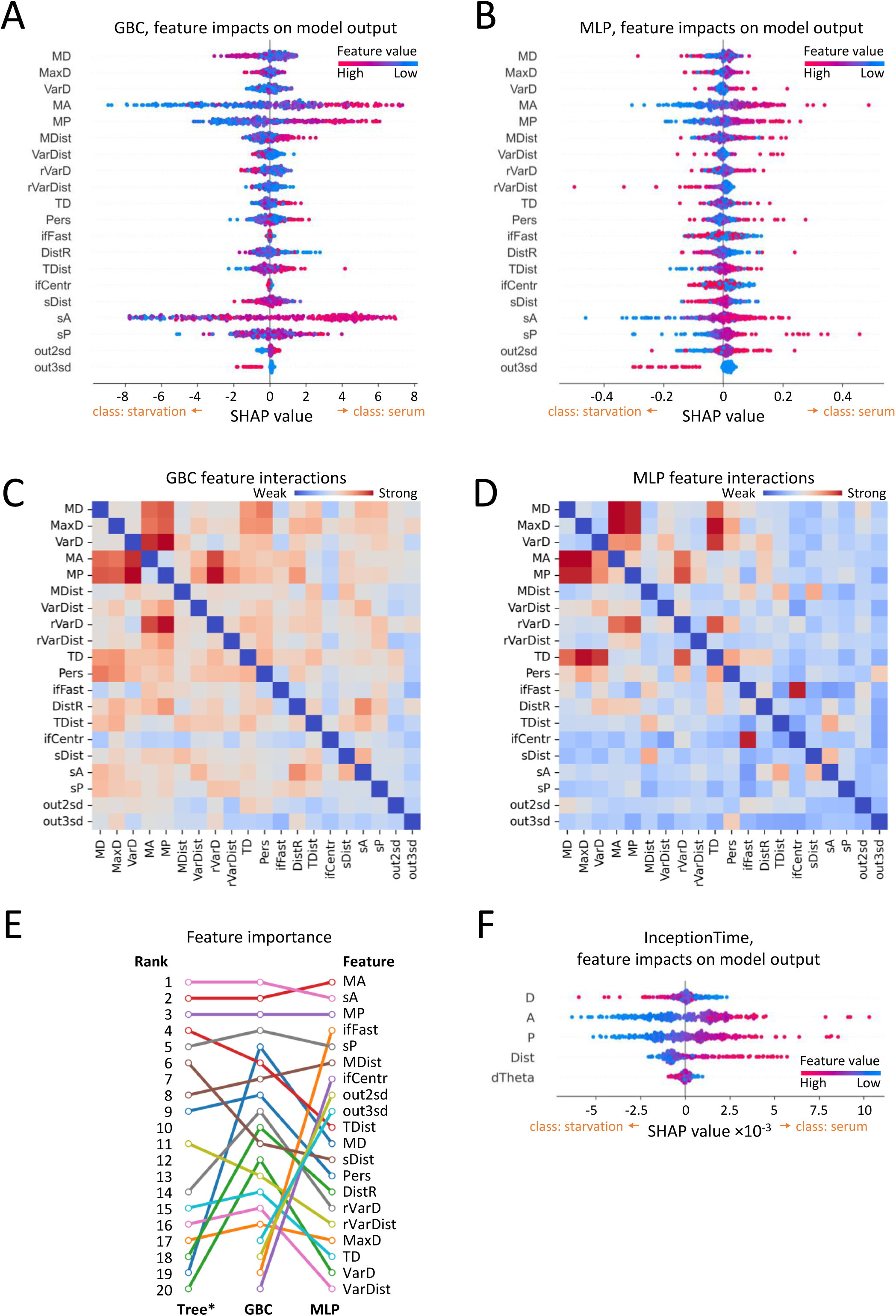
Explaining machine learning models’ decision using Shapley additive explanations (SHAP). **A,B** Global summary of SHAP showing feature (see Table 1) impacts towards one of the classes, high serum or starvation, for gradient boosting classifier (GBC) (**A**) and multilayer perceptron (MLP) (**B**).Feature values are color-mapped from blue (low) to red (high). **C,D** Heatmap of feature interactions depicting the components of synergistic effect of features, after accounting for the direct impact. Interactions from GBC seem to be more uniform and show less usage of binary and discrete features (**C**), compared to the interactions from MLP (**D**). **E**, Change in feature importance rank with the growth of model complexity from the baseline single level decision tree (designated as Tree) to GBC and MLP. Binary and discrete features are not shown for the Tree set, as the model is trained on a single feature at a time to estimate baseline performance, thus, feature interactions are not used by definition. **F**, Global summary of SHAP showing feature impacts towards class serum or starvation, InceptionTime.

Since chromosome 1 in U2OS cells predominantly has two homologs in each nucleus, information gain from *ifFast* and *ifCentr* cannot be explained by the impact of the feature value itself but rather implies the influence of these features on other features’ SHAP values. Fortunately, SHAP enables estimation of interaction components separated from direct impact of the feature value itself (Molnar, 2019). Indeed, the *ifFast* and *ifCentr* did not have any noteworthy hits on GBC interactions map, while showed strong interaction with each other on MLP (Fig 3C,D). MLP feature interactions map in general seemed less noisy, highlighting also the interactions between speed and morphology and total locus displacement and maximal displacement within the timelapse (Fig 3C,D). Altogether, the SHAP explanations and the interactions derived from SHAP analysis allowed us to better navigate in the feature space of the dataset, narrowing further analysis to the interpretation of important patterns, as described below.

### Nuclear morphology during serum starvation

The explanation of models trained on chromosome 1 tracking data to distinguish high serum condition from serum starvation showed a significant impact of morphological features on classifier decision. Nuclear area (*MA*) and the trend of its change (*sA*) demonstrated highest importance as indicated by SHAP analysis of both GBC and MLP (Fig. 3E). Nuclear area also had highest mean absolute SHAP value in InceptionTime predictions (Fig. 3F). Indeed, a change in the skewness of nuclear area distribution was observed under starvation (Fig. 4A) with the detected decrease in the mean value of approximately 50 squared microns, which represents more than 10% of the average nuclear area (Fig. 4B, Supplementary Figure 4A). Importantly, this change alone cannot explain the radial rearrangement of chromosome 1, as we did not detect any change in telomere distance to the nuclear edge (Fig. 1F). The nuclear area also becomes more dynamic under serum starvation (Fig. 4C), and the changes in both morphological features were detectable already within the first 20 min after the medium change (Fig. 4B,C). Notably, nuclei with an area below 250 squared microns rarely demonstrated further decrease in their area (Fig. 4D), perhaps due to the area decrease occurring prior to timelapse recording. Since the length of the nuclear perimeter (MP), similar to the area (MA), decreased in starvation (Supplementary Figure 4B), we wondered whether this shrinkage led to a change in circularity, similarly as reported for apoptotic cells (Toné *et al*., 2007; Helmy and Adel, 2012). In our dataset, the changes in both circularity and the correlation between area and perimeter were negligible (Fig. 4E, Supplementary Figure 4C), confirming the different nature of serum starvation-driven changes in nuclear morphology compared to apoptosis.

**Figure 4.**
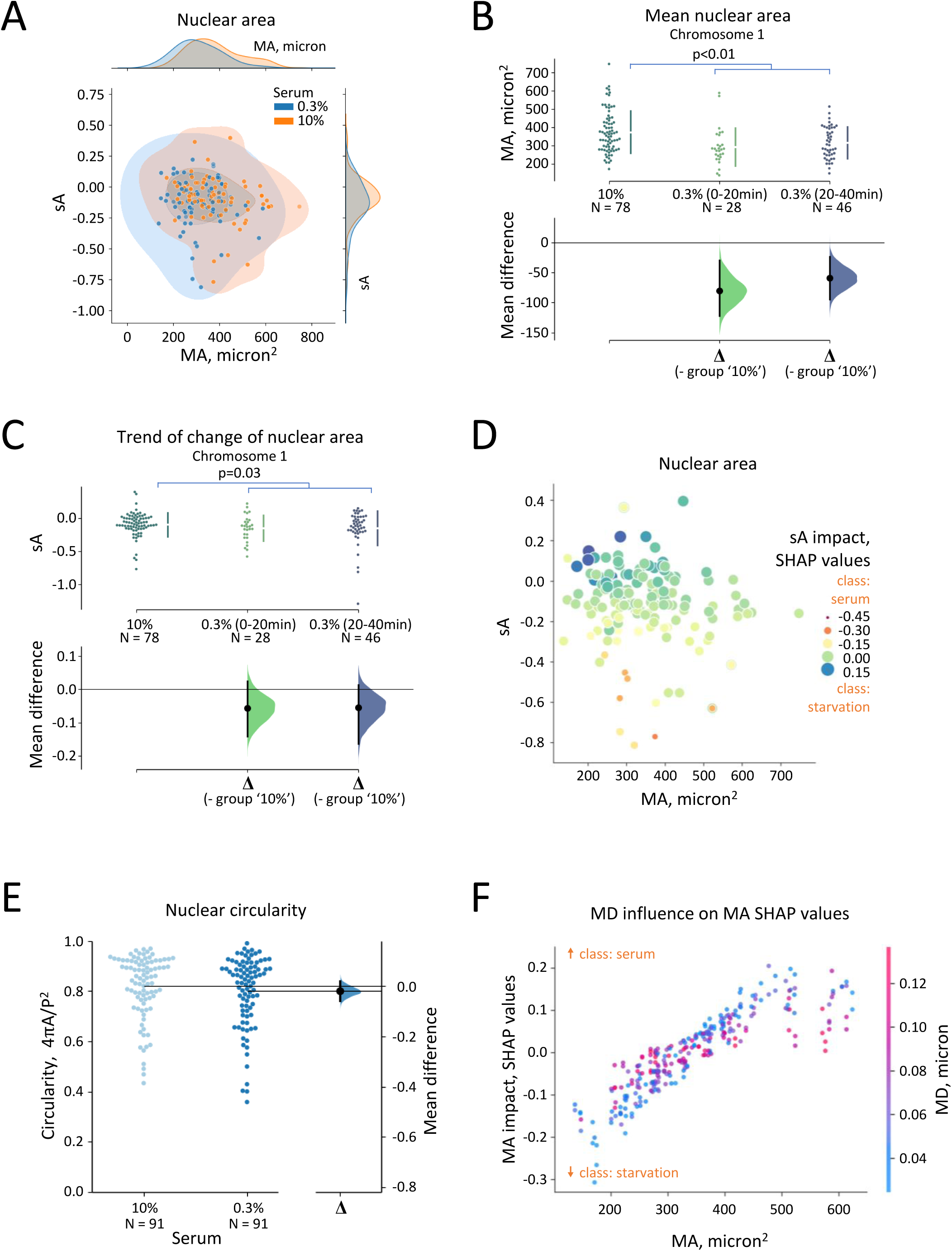
Changes in nuclear morphology upon serum starvation. **A**, Nuclear area (MA) and its dynamics (sA) in high serum medium (10%) and under starvation (0,3%). The marginal distributions show corresponding kernel density estimations. **B-E**, Changes in nuclear morphology, particularly, area (**B**) area dynamics (**C,D**) and circularity. Statistical significance with Mann-Whitney U rank test. (**E**). Each dot in the swarmplot represents an individual nucleus; number (N) of analyzed nucleus indicated below the swarmplot. Panels on the right represent the distributions of bootstrapped delta mean values for datasets. Error bars show 95% confidence intervals. **D**, Impact of nuclear area dynamics (as SHAP values, shown in color) on model decision depending on nuclear area and dynamics of its change. **F**, Locus speed as mean displacement (MD) affects Shapley values for nuclear area, which is clear from vertical color gradient, where the locus speed is color-mapped, see scale on the right. MLP SHAP explanation; outliers (less than 2% of data) are not shown for clarity.

Further, the SHAP explanation of both GBC and MLP revealed interactions between speed-related features, such as mean (MD) and maximal (MaxD) locus displacement and morphological features, namely *MA* and *MP* (Fig. 3C and more prominent in Fig. 3D). Upon closer examination, we noticed that high mean displacement values shifted the area’s impact on model decisions towards SHAP values around zero (Fig. 4F). This was also true for GBC and InceptionTime explanation (Supplementary Figure 4D,F), thus showing that the predictive value generated by nuclear area is rather contextual, e.g. its importance is high in a subgroup of low motility loci. In summary, our results demonstrate serum starvation induces a global change in nuclear morphology.

### Unique characteristics of chromosome 1 loci movement under starvation

To achieve the most comprehensive integration of morphology data and chromosome 1 loci motility, we next augmented the basic raw displacement data with prior knowledge-based engineered features. Initially, no obvious change in the speed of chromosome 1 loci, as measured by mean displacement (*MD*), was observed under starvation (Fig. 5A). However, telomeres seemed to display a trend of decreased motility (Fig. 5B), which could be explained by the decrease in nuclear volume (Fig. 4B), possibly resulting in increased chromatin density. At the same time, the total distance traveled by chromosome 1 loci within the timelapse (*TD*) grew in parallel with speed (Fig. 5A,C), while no growth in total distance was observed for the telomeres (Fig. 5D). Thus, chromosome 1 loci seem to retain their motility level, while telomeres, on average, seem to slow down under starvation. Interestingly, while both nuclear morphology (Fig. 4B,C) and telomere dynamics (Fig. 5B) responded already during the first 20 minutes of serum starvation, the changes for chromosome 1 were observed later (Fig 5A,C), further highlighting the specific response of chromosome 1 loci for this particular signal.

**Figure 5.**
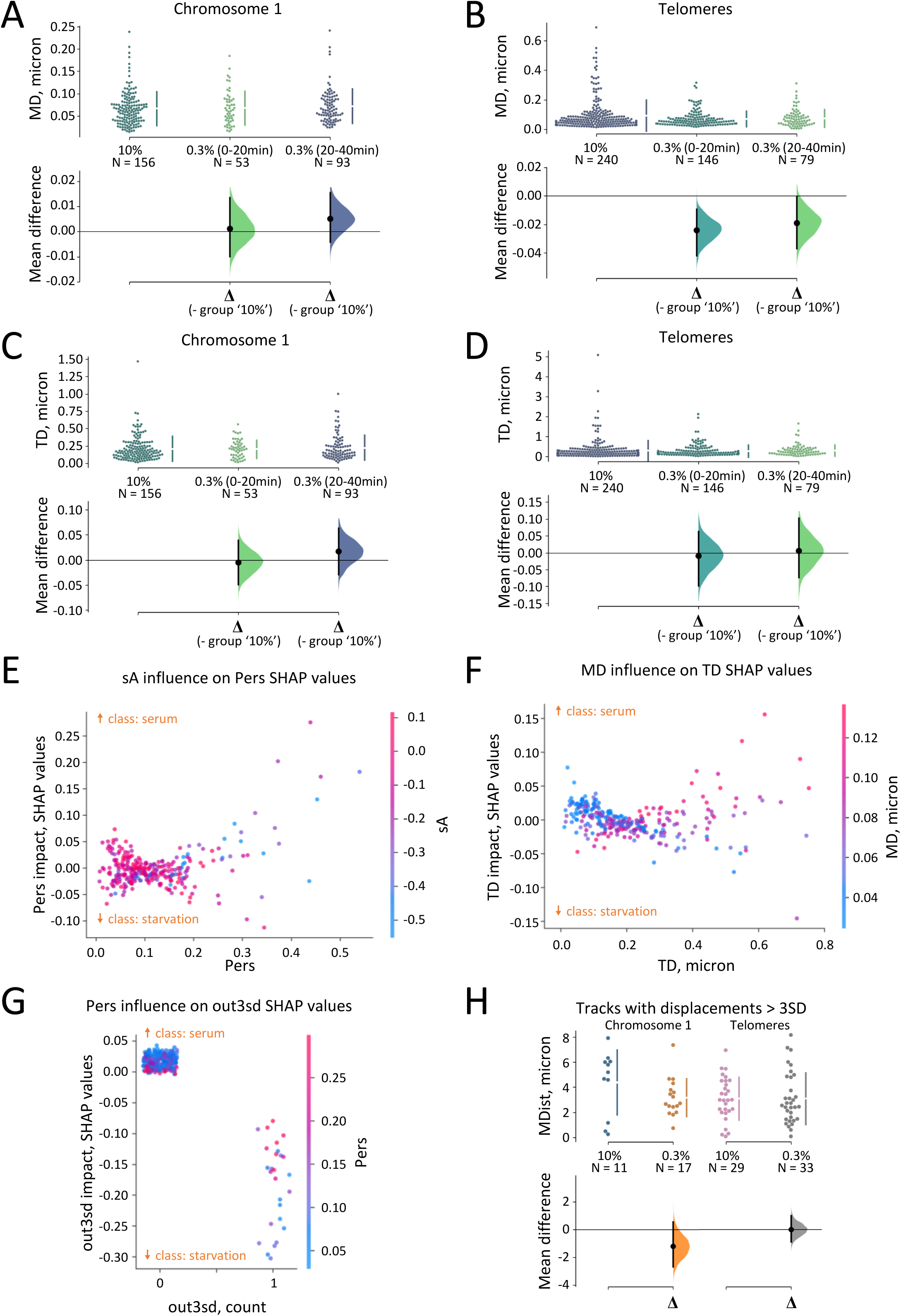
Specific features of chromosome 1 dynamics. **A-D**, Changes in loci motility as measured by mean displacement (*MD*) and total distance traveled (*TD*). Mean loci displacement did not change for chromosome 1 under starvation (**A**), but showed a decrease for telomeres (**B**). Total distance traveled by a locus for chromosome 1 grew insignificantly, in a very similar manner to its mean displacement (**C**), while for telomeres it stays stable despite the decrease in speed (**D**). Each dot in the swarmplot represents an individual locus; number (N) of analyzed loci indicated below the swarmplot. Panels on the bottom represent the distributions of bootstrapped delta mean values for datasets. Error bars show 95% confidence intervals. **E-G**, SHAP analysis of features describing persistence and directionality of loci movement. **E**, dynamics of the change of nuclear area (*sA*) affects the persistence (*Pers*) SHAP values. **F**, Locus mean displacement (*MD*) affects the SHAP values calculated for total traveled distance (*TD*). **G,** Persistence (*Pers*) is negatively correlated with SHAP values of feature which detects long spontaneous displacements (*out3sd*). **H,** Long spontaneous displacements (*out3sd* = 1) of chromosome 1 loci are more likely to occur in short distances from nuclear periphery under serum starvation, data shown as in A-D.

To further characterize chromosome 1 dynamics, we next focused on persistence, calculated as the ratio of total distance traveled (*TD*) to the trajectory length (*MD*\*n_frames_). The initial assumption, based on FISH-based studies, that long-range persistent movement would be detectable under starvation did not hold, as persistence did not correlate with persistence SHAP values (Fig. 3A,B, Fig. 5E). At the same time, SHAP values for total distance (*TD*) depended on loci speed in a non-linear way, with low-speed loci (TD < 0.2) having close to zero SHAP value (Fig. 5F). In the high TD range, generally slow-moving loci are associated with negative SHAP values (Fig. 5F), thereby influencing the model’s decision towards the ’starvation’ class. This dependence suggests that speed of locus movement and its directionality do interact in a meaningful way that is specific to chromosome 1 loci. However, this interaction cannot be reduced to the simple measure of persistence.

Remarkably, the presence of unusually long (*MD*+3σ) displacements in chromosome 1 track (feature named *out3sd*), demonstrated a considerable and uniform impact on MLP decision, directed towards starvation class (Fig 3A,B, Fig. 5G). The SHAP values of *out3sd* were negatively correlated with persistence (Fig. 5G), suggesting that *out3sd*-outliers do not just follow the slow unidirectional movement. Instead, the loci seem to undergo random walk with occasional fast relocalizations. Moreover, under starvation, these long displacements for chromosome 1 loci, but not for telomeres, were detected with higher probability in the proximity of the nuclear envelope (Fig. 5H). Thus, the ability to maintain dynamics in the smaller nuclei (Fig. 5A) alongside intermittent ’jumps’ (Fig. 5G) especially near the nuclear envelope (Fig. 5H) appear to be hallmarks of chromosome 1 dynamics in a low serum environment.

### Radial organization and differential behavior of homologous chromosomes

Being intrigued by the higher occurrence of long displacement outliers close to the nuclear periphery (Fig. 5H), we analyzed in more detail the set of features related to radial localization of loci, namely *DistR* and *TDist*. *DistR* is the absolute difference between the initial and final radial positions within the timelapse, which does not show the direction, towards the center or the periphery, of the movement. *DistR* exhibited a slight increase in starvation for chromosome 1 loci, while demonstrating a pronounced increase for telomeres (Fig. 6B). In contrast, *TDist* is the total radial distance traveled by a locus, and shows the direction of relocalization. For chromosome 1 loci, the distribution of raw values for *TDist* skewed downwards in serum starvation (Fig. 6C), supporting the relocalization towards the nuclear periphery. The distribution for change of *TDist* in telomeres was much wider, with the mean value closer to zero, while raw values for telomeres were distributed symmetrically (Fig. 6C). Thus, also these features support the chromosome-specificity of radial reorganization.

**Figure 6.**
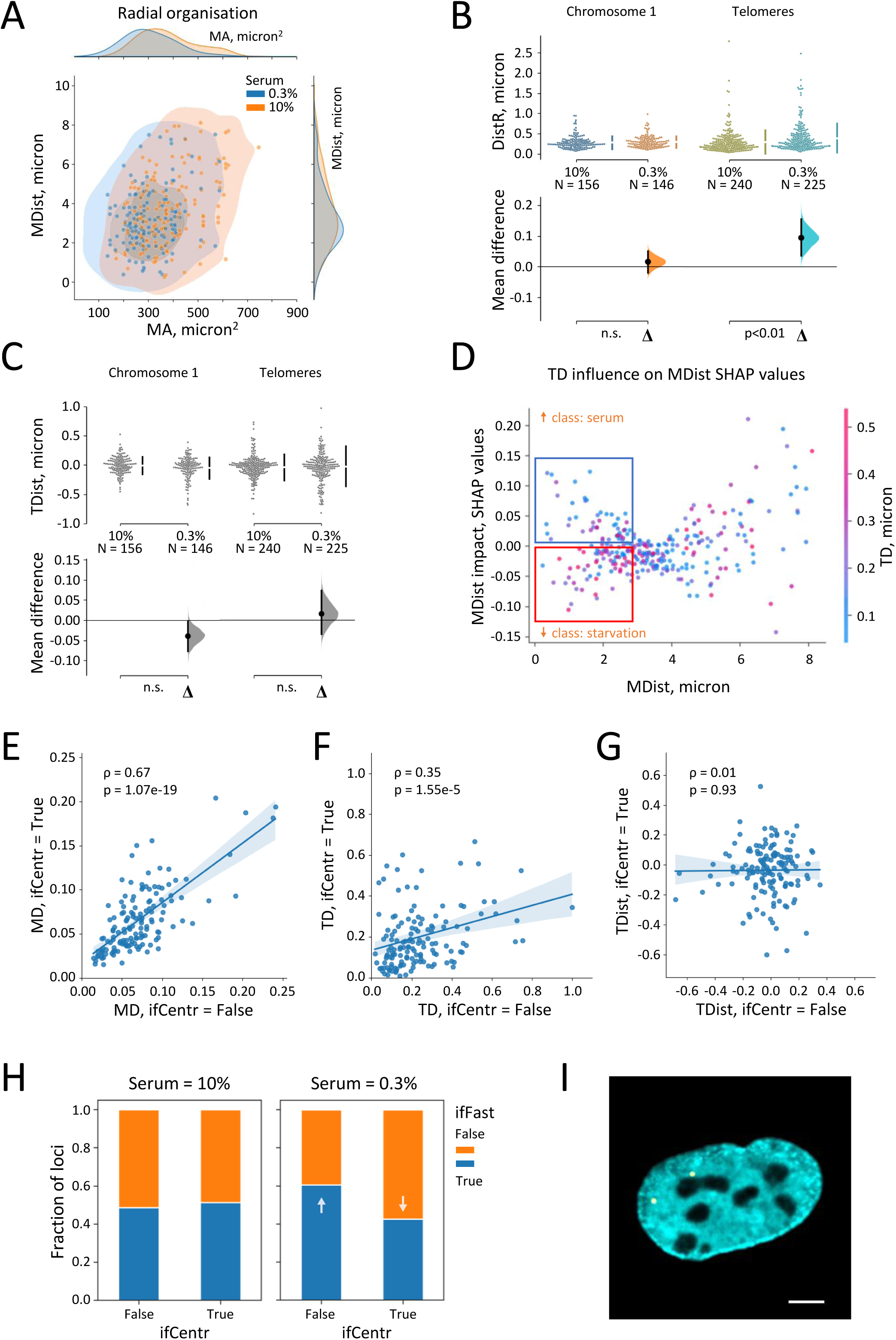
Distinct dynamic behavior of chromosome 1 homologs upon serum starvation A-D, Chromosome 1 radial organization as explored using time series data analysis. **A**, Nuclear area (MA) and mean locus distance to nuclear edge (MDist) in high serum medium and under starvation. The marginal distributions show corresponding kernel density estimations. **B,C** Timelapse-derived data confirm chromatin radial reorganization under serum starvation. The dynamic range of radial movement (DistR) changed insignificantly for chromosome 1, but showed substantial growth in telomeres (**B**). Total length of radial relocalization (TDist), showing not only scale but also the direction of relocalization, decreased in chromosome 1 but had only insignificant change in telomeres (**C**) Statistical significance (**B,C**) with Mann-Whitney U rank test. **D**, Total distance traveled by locus (TD) affects mean distance to edge (MDist) SHAP values. **E-I,** Differential behavior of Chromosome 1 homologs. **E-G**, Correlation of mean loci displacement (MD) (**E**), total distance traveled by locus (TD) (**F**), and total radial displacement (TDist) (**G**) between chromosome 1 homologs. **H**, Relationship between categorical features subgrouping chromosome 1 homologs based on speed (ifFast) and radial position (ifCentr). **I**, Representative image of chromosome 1 loci (in yellow), where one of the homologs is associated with a nucleoli-like subcompartment. The LaminA channel is shown in cyan. Scale bar 6 microns.

In line with the observations on radial reorganization (Fig. 1F, Fig. 6C), SHAP dependence plot of the ‘static’ distance to edge, *MDist*, in GBC showed positive correlation between *MDist* and its SHAP (Fig. 3A). While MLP explanation did not show similar clear correlation (Fig. 3B), its butterfly-like shape had a narrowing distribution of SHAP-values at around 3-4 microns from the edge (Fig. 6D). This narrow distribution indicates an area of less predictive power and alludes to the division of the dataset to two separate subclasses - peripheral and central. On the *MDist* SHAP dependence plot based on MLP decisions (Fig. 6D), high total displacement (*TD)* in peripheral subclass pushes the prediction towards starvation class, in line with higher chance to see the out3sd outliers in the same region (Fig. 5H). These observations suggest the relevance of subgrouping the chromosome 1 homologs based on radial localization.

To ascertain the influence of radial localization on the differential behavior of chromosome 1 homologs, the loci were grouped based on two factors: speed (*ifFast*) and proximity to the periphery (*ifCentr*). In a primary screening, locus speed (*MD*), total traveled distance (*TD*) and total radial displacement (*TDist*) were correlated between homologs regardless of serum starvation status (Fig. 6E-G). Significant positive correlation between homologs’ speed (Fig. 6E) and total traveled distance (Fig. 6F) was detected, likely showing that those features largely depend on the general degree of constraint in the particular nucleus. If the radial reorganization would be caused by non-specific deformation of the nucleus, the total radial distance traveled by a locus (*TDist*) should be correlated for both homologs. However, our observation shows no correlation (Fig. 6G), indicating that general structural distortion does not affect radial reorganization. This suggests that certain chromosome-specific effects, which may independently affect homologs, could orchestrate radial reorganization during serum starvation.

Curiously, the MLP SHAP explanation highlighted the interaction between the features generating homolog subgroups, *ifFast* and *ifCentr* (Fig. 3D). To further examine this finding, we investigated how the *ifFast* label is distributed between the central and peripheral homologs in the different conditions. In the high serum condition, *ifFast* and *ifCentr* are well-balanced, indicating that the peripheral locus has an equal probability of being faster or slower compared to its homolog (Fig. 6H, left). However, under starvation, there is a shift in this distribution, associating peripheral loci more frequently with the ifFast label (Fig. 6H, right). This indirectly supports (i) the findings from the MDist SHAP analysis, which showed that high total displacement (*TD*) is not predictive in central (*MDist* > 3 microns) loci (Fig. 6D) and (ii) a higher likelihood of detecting fast spontaneous movements closer to the periphery under starvation (Fig. 5H). Overall, this characterization indicates that peripheral loci are more mobile and responsive to stimuli. Intuitively, this may be related to the frequent detection of one of the homologs, often the central one, associated with a subcompartment resembling nucleoli (Fig. 6I). Thus, in serum starvation, chromosome 1 homologs display differential behavior, with peripheral loci demonstrating greater mobility and responsiveness to stimuli.

## DISCUSSION

Chromatin is dynamic at all length scales, and methods for sequence-specific labeling of chromatin loci are opening new avenues for studying the underlying molecular mechanisms with live imaging techniques. As the dynamics of a specific locus are influenced by the overall chromatin organization, proximity to nuclear landmarks as well as the general nuclear morphology, extracting the relevant patterns from multidimensional imaging data becomes a challenge. Here we have combined CRISPR-Sirius-based labeling of specific chromatin loci for live imaging with machine-learning assisted dataset exploration followed by model explanation with SHAP. This approach uncovered the specific features of chromosome 1 loci dynamics during serum starvation and is widely applicable to dissect mechanistic insights from many different types of experimental data.

Given that ML models vary in efficiency and might highlight different aspects of the dataset under study, a comprehensive exploratory analysis normally includes testing a set of models (Xu and Jackson, 2019; Deng *et al*., 2021). In our hands, while all the trained models were superior to the baseline (single-level decision tree), MLP, trained on the prior knowledge-based engineered features aggregated over time-axis, had the highest accuracy, presumably extracting the most information from the available dataset (Fig. 2E,F,G). Despite drastically different architecture, in many cases the trained models generated notably similar patterns of feature importances. For example, locus distance to nuclear periphery was positively correlated with prediction of class ‘serum’ by both GBC and InceptionTime (*MDist* and *Dist*, Fig.3A,F). Another example includes intricate interaction between nuclear morphology and locus speed. For all the used models, a large nuclear area was associated with class ‘serum’, which is evident from positive correlation between area and area SHAP values (Supplementary Figure 4D-F). At the same time, for the subgroup of fast loci (high D, MD) this correlation was lower, while slow loci were characterized by highest absolute SHAP values. In the GBC explanation, this pattern was least clear, but still recognizable; InceptionTime showed SHAP values distribution similar to MLP, with even stronger correlation (Supplementary Figure 4F). Overall, this similarity suggests that all the models were able to create the aligned abstractions, accurately approximating the underlying data.

On the other hand, despite general trends being captured by all tested models, certain patterns were specifically highlighted by some of them. Notably, displacement length (*D*) clearly correlated with the starvation class prediction only for InceptionTime. Since this model preprocessed raw time-resolved data by ‘scanning’ it with one-dimensional convolutions, one possible explanation for this observation is that InceptionTime modules effectively extracted specific events or patterns from the time series, analogous to how *out3sd* events were manually extracted to train GBC.

Features describing nuclear morphology showed high importance for decisions of all tested classifiers (Fig. 3E), highlighting the profound impact of serum starvation on the cell. Serum starvation is a signal for the cells to become quiescent and exit the cell cycle. It thus affects numerous signaling pathways, and genomic functions from epigenetics to gene expression (Pirkmajer and Chibalin, 2011). It is important to note that here we focused on the very immediate response to starvation and monitor nuclear and chromatin response only during the first hour after changing the media to low serum. The functional implications of chromosome repositioning upon serum starvation are not obvious, but it is tempting to speculate that repositioning of chromosomes to new nuclear environments optimizes their function in the changing environment. For example, the differences in chromatin dynamics observed within the first 20 minutes compared to 20-40 minutes of starvation (Fig. 5A,B) could indicate early and late changes in gene transcription. Further studies correlating dynamics with functional outputs of the genome are required to elucidate these connections. Previous studies have shown that almost half of the chromosomes displayed altered radial localization in proliferating compared to quiescent or serum starved cells (Mehta *et al*., 2010). Indeed, we found that although telomeres in general became less mobile upon serum starvation (Fig. 5B), perhaps due to increased chromatin density upon decreased nuclear volume (Fig. 4B), they displayed increased radial movement without preferential orientation towards nuclear center or periphery (Fig. 6C). This may reflect the overall radial movement of several chromosomes, which could hinder the movement in the lateral dimension. Previous studies with CRISPR-Sirius had indicated cell-cycle dependent chromatin dynamics, with the chromatin movement increasing from early to late G1 phase of the cell cycle, and then decreasing during the S phase (Ma *et al*., 2019). Here we have restricted our analysis to G1-early S phase of the cell cycle, since we analyzed only cells displaying two “dots” of chromosome 1 loci. In the future, it would be interesting to analyze whether the cell cycle state influences the chromatin response to serum withdrawal. Nevertheless, the fact that we detected radial reorganization of chromosome 1 loci also when analyzing the same cell in the presence and absence of serum (Fig. 1G), indicates that the cell cycle stage is not sufficient to explain the changes in chromatin dynamics.

The overall change in nuclear morphology (Fig. 4A-D) and chromatin dynamics (Fig. 5B) do not explain the radial relocalization of chromosome 1 loci (Fig. 1F,G) in response to serum starvation. Instead, chromosome 1 loci display distinct dynamic behavior, which is characterized by maintaining the dynamics (Fig. 5A) in the otherwise constrained environment (Fig. 5B), and “long-jumps” (Fig. 5G), especially near the nuclear periphery (Fig. 5H). One example demonstrating the power of SHAP is the interpretation of persistence in locus movement. Since the FISH-based experiments had suggested a role for an actomyosin motor in chromosome repositioning (Mehta *et al*., 2010), we expected to see evidence of active, directed movement upon serum starvation, similarly as reported, for example, for heat-shock induced repositioning of the HSP70 transgene (Khanna *et al*., 2014) or movement of heterochromatin breaks in fly cells (Caridi *et al*., 2018). However, there was no noticeable gain in predictive power from persistence, calculated as *TD/MD*\*n_frames_ in any of the trained models. Nevertheless, *TD* had a pronounced interaction component in its impact on the model’s decision, highlighting its interaction with speed-related movement characteristics (Fig. 3D). Further analysis identified a subgroup of loci that are characterized by high total displacement but low mean displacement under serum starved conditions (Fig. 5F), and suggested an association with another feature, namely *out3sd*, which indicates the presence of long displacement outliers within the captured timelapse. In the future, improving the temporal resolution of imaging could provide valuable insights into the nature of the observed fast intermittent movement and potentially prove the role of active transport, perhaps mediated by a nuclear actomyosin motor. Another intriguing question is how the directionality of relocalization is established for specific chromatin loci. It would, for example, be interesting to analyze the dynamics of further chromatin loci, such as those moving towards the center of the nucleus.

Majority of the reports on radial chromosome organization neglect the difference in behavior between homologs, operating either with averaged fluorescent signal measured within a radial mask (Mehta *et al*., 2010), mean centroid position (Mehta *et al*., 2021), or working with haploid cell lines like HAP1 (Girelli *et al*., 2020). However, there is a body of data suggesting that a considerable number of genes, apart from inactivated X chromosome, exhibit monoallelic expression (Gimelbrant *et al*., 2007). Based on these data and measurements of inter-homologue distances, it has been suggested that homologous chromosome territories are positioned in a non-random manner to allow independent regulation, potentially involving the participation of active mechanisms for functional interchromosomal interactions (Heride *et al*., 2010). In this study we report the differential response of homologous loci to serum starvation. Peripheral loci demonstrated longer total displacement when compared to central ones (*TD*, Fig. 6F), and total radial relocalization, *TDist*, did not correlate between homologs (Fig. 6G). Moreover, the proportion of faster peripheral loci increased under starvation (Fig. 6H). One explanation to the differential behavior of chromosomes may be anchorage to specific nuclear substructures. Indeed, we often observed one of the homolog loci in association with structures resembling nucleoli (Fig. 6I), which represent a heterochromatin-associated subcompartment (Bizhanova and Kaufman, 2020) similarly as the nuclear periphery. In addition, differences in transcriptional or chromatin states between the homologs could play a role. Thus, starvation-driven change in chromatin dynamics can be specific not only to chromosomes, but also to specific homologs.

In general, the explorative approach proposed in this study is conceptually aligned with diffusional fingerprinting as described by Pinholt et al. (2021). While both techniques share the idea of using the enriched feature space to analyze particle tracking data, there are several key distinctions. By design, diffusional fingerprinting employs feature engineering primarily focused on particle biophysics, with the intention to provide generalizable model applicable across diverse types of diffusional behavior. In contrast, we designed the feature set to specifically characterize inherently constrained chromatin movement, while also integrating the relevant data on nuclear morphology. Additionally, the two methods differ in their explainability frameworks. While diffusional fingerprinting utilizes linear discriminant analysis to extract mechanistic insights into particle behavior, SHAP, used in our study, enables greater flexibility when used with complex ML models. The observed relationships between radial organization, speed, total displacement, and directionality suggest that within the analyzed feature space, the machine learning models were able to identify patterns that are more informative about the experimental conditions under study than any of the manually engineered features based on prior knowledge by itself. We believe that the reported dataset exploration technique offers a powerful tool for effectively mining complex patterns within datasets in both basic biology and biomedicine.

## MATERIALS AND METHODS

### Plasmids and guide RNA

CRISPR-Sirius system plasmids, namely pHAGE-TO-dCas9 (Addgene ID 75381), pHAGE-EFS-PCP-GFPnls (Addgene ID 121938), pLH-sgRNA-Sirius-8xPP7 (Addgene ID 121940) were kindly provided by Pederson lab (Ma et al., 2018). sgRNA was cloned into pLH-sgRNA as described in (Ma et al., 2016) using Stbl3 competent cells. sgRNA sequences:

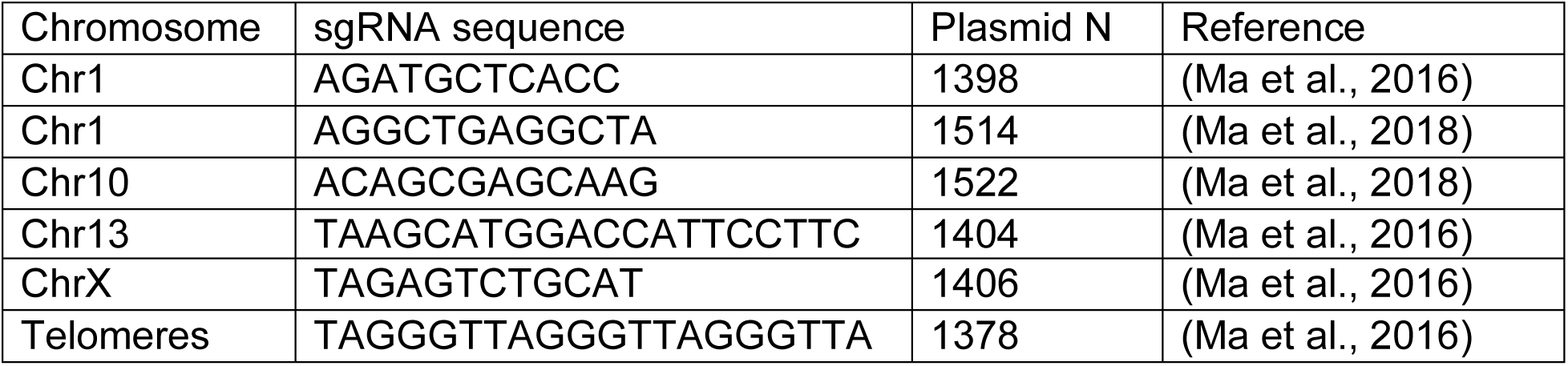

Chromosome localization of the target sequences is shown on Supplementary Figure S1A. Guide RNAs specific to CTGGGTCCCTGT (Chr10, plasmid N1521), AGGCTCCAGACC (Chr10, Plasmid N1520), ACCATTCCTTC (Chr13, Plasmid N1403), and TGCTCCACCCAA (Chr1, Plasmid N1515) were tested but corresponding loci were not visible in our experiments, thus were not used in this study.

LaminA-mCherry construct was generated by swapping EGFP to mCherry in pEGFP-LaminA vector. Plasmids created for this study are available upon request.

### Live cell imaging

For live cell imaging, human U2-OS cells, a kind gift from Lappalainen-lab (University of Helsinki) and regularly tested for mycoplasma contamination, were cultured routinely in DMEM medium supplemented with 10% FBS (GIBCO), 100 Units/ml penicillin, and 0.1 mg/ml streptomycin (Pen-Strep, GIBCO), then trypsinized and transferred to 35mm glass bottom Mattek dish following transfection with Lipofectamine 3000 (Thermofisher) according to manufacturer’s protocol. 200 ng of pHAGE-TO-dCas9, 40 ng of pHAGE-EFS-PCP-GFPnls, 1000 ng of pLH-sgRNA-Sirius-8xPP7 and 150 ng of LaminA-mCherry plasmids were used per 35 mm dish. Live cell imaging was performed 24-36 h after transfection in FluoroBrite DMEM (Thermofisher). First, cells were imaged in 10% serum (FBS, GIBCO), and then medium changed to 0.3% serum. Imaging was performed using an Intelligent Imaging Innovations (3i) Marianas Inverted Spinning Disk (Yokogawa CSU-X1 M1 5,000 rpm) microscope, equipped with Andor Neo sCMOS camera and solid-state lasers (488 nm/150 mW, 561 nm/50 mW). Cells were imaged using a 63x/1.2 W C-Apochromat Corr WD=0.28 M27 (Zeiss). Data is from at least three biological replicates, with the number of loci analyzed indicated in the figures. For the experiments where the same cells were imaged before and after the medium change (paired data), XY coordinates were stored for the set of cells and the medium was changed on the stage. The CO2 level and temperature were controlled using the environmental chamber and CO2 mixer with active humidity control (Life Imaging Services). For each cell, a Z-stack of 5 sections was captured, with 1 micron step size, 2 min total timelapse duration, 0.25 frames per second. Since performing statical error control on 3i Marianas was not possible for technical reasons, imaging of fixed cells was done on Aurox Clarity Zeiss, which shared the microscope frame, stage, and optics including objective with the Marianas, to ensure comparability of the results. Images were analyzed using the same localization detection algorithm (Crocker and Grier, 1996) as implemented in *trackpy* library (version 0.5.0) used for the rest of the data (see below).

### Image segmentation, loci tracking and data pre-processing

Image format conversion from proprietary .sld (3i) to .ome.tiff (OME) was done using Bio-Formats plugin (Linkert et al., 2010) before the z-projection as a part of custom-made script in ImageJ, 1.51 (Schneider *et al*., 2012). Next, files and metadata were read in python using *apeer_ometiff_library*. Nuclei were segmented using multi-Otsu and morphology toolbox from *scikit-image*, based on LaminA-mCherry channel, to get nuclear morphology characteristics, namely nuclear area and perimeter. Nuclear circularity was approximated as 4***π*A** / **P**^2^, where **A** and **P** stand for area and perimeter correspondingly. Rigid body registration (*pystackreg*) was used to compensate for the nucleus translation and rotation between frames. Chromosome loci visualized by CRISPR-Sirius were detected and tracked by the *trackpy* (Crocker and Grier, 1996), to acquire raw data on loci displacements in between frames. The binary masks of nuclear perimeter and xy coordinates of chromosome loci were used for mapping the loci distance to nuclear rim using *numpy*. Finally, the data collected in tracking experiments was aggregated, cleaned and further used for prior knowledge-based feature engineering mainly using *pandas, numpy* and *scipy.stats*. For more details of image analysis please see the code, link provided in the Data Availability section.

### Machine learning on tabular data

#### Pipeline architecture

The core idea behind ML-assisted dataset exploration involves training ML models on tracking data using experimental conditions as labels. The Shapley values-based approach is then employed to explain the models and uncover higher-level descriptive patterns used by the models for classification. An ensemble of gradient boosted decision trees (*scikit-learn*, designated GBC) and a small 400k parameter neural network consisting of seven fully-connected layers (*keras*, designated in text as MLP) were chosen due to robustness, computational lightweightness and accessibility regardless of operators level of technical knowledge.

#### Baseline

The classes in the dataset under study were almost perfectly balanced, thus threshold for predictive power set at ZeroR = 0.52. To evaluate the ability of GBC and MLP to infer relatively complex patterns from the available data, we established an additional baseline by measuring the performance of a single-level decision tree using the same evaluation scheme employed for GBC and MLP.

#### Hyperparameters screen and performance evaluation

A hyperparameter search was conducted for GBC, exploring the range of learning rates and maximal tree depths; for the MLP architecture, the depth and number of neurons in dense layers were manually adjusted for tuning. The optimal configurations were further evaluated using repeated StratifiedGroupKFold cross-validation, with the groups defined by association to a specific nucleus, therefore preventing the data leakage due to shared morphology features. Data preprocessing, grid search and cross-validation were done in *scikit-learn* environment. For InceptionTime, the number of inception modules (affecting the overall model size) and the kernel sizes of the modules’ 1D convolution layers (the time window lengths) were altered for hyperparameter tuning.

#### SHAP explanation

Shapley values-based explanation (Lundberg and Lee, 2017) was done in parallel with cross-validation. Calculated values were aggregated and accumulated through cross validation repeats, taking advantage of additivity, an intrinsic feature of Shapley values. The approximate feature interaction index values were calculated using the *potential_interactions* function from *shap* utils, with the final interaction matrix (**A**_si_) being calculated as entrywise sum of the output from *potential_interactions*, designated **A**_i_ and its transposed version (**A**_i_^t^), divided by two (scalar multiplication), as in **A**_si_ = 0.5(**A**_i_+**A**_i_^t^)

#### Data visualization

Data was visualized using *matplotlib*, *seaborn* and *dabest* (Ho *et al*., 2019). SHAP visualizations were done using a built-in toolbox from the *shap* library. Descriptive statistics data are shown on figures as kernel density estimation, confidence intervals and bootstrapped changes distributions. Statistical significance was tested with Mann-Whitney U rank test (independent samples, two-sided) or t-test (paired, two-tailed), as indicated. See also Supplementary table 1 for inferential statistics with two-sided Mann-Whitney (MW) U-test.

## DATA AVAILABILITY

The code used in this study is available at https://github.com/redchuk/chromatin-tracking-exploration

## AUTHOR CONTRIBUTIONS

MKV and TR designed the study. ON cloned and tested the sgRNA constructs for CRISPR-Sirius; TR, APP and ON performed the imaging. TR and HJ wrote the code and performed the data analysis. LP helped to design machine learning experiments and performed code review. MKV and TR did the biological interpretation and wrote the manuscript. All authors have seen and approved the manuscript.

## ACKNOWLEDGMENTS

Microscopy was performed at Light Microscopy Unit (Institute of Biotechnology, HiLIFE) and authors are grateful to the staff for technical assistance. The authors wish to acknowledge Scientific Computing Services (HPC, University of Helsinki) and CSC – IT Center for Science, Finland, for computational resources. TR would like to thank Rustem Kasymov (Noice Entertainment Oy had no role in the study), Ilya Belevich (EMU, UH), Redchuk AS, and members of Viikki Bioinformatics club (UH) for the insightful discussions. This work was supported by Sigrid Juselius, Cancer and Jane and Aatos Erkko foundations as well as Academy of Finland grants 338281 and 330254 to MKV. Open access funded by Helsinki University Library.

## Supplementary Material

**Supplementary figure 1.**
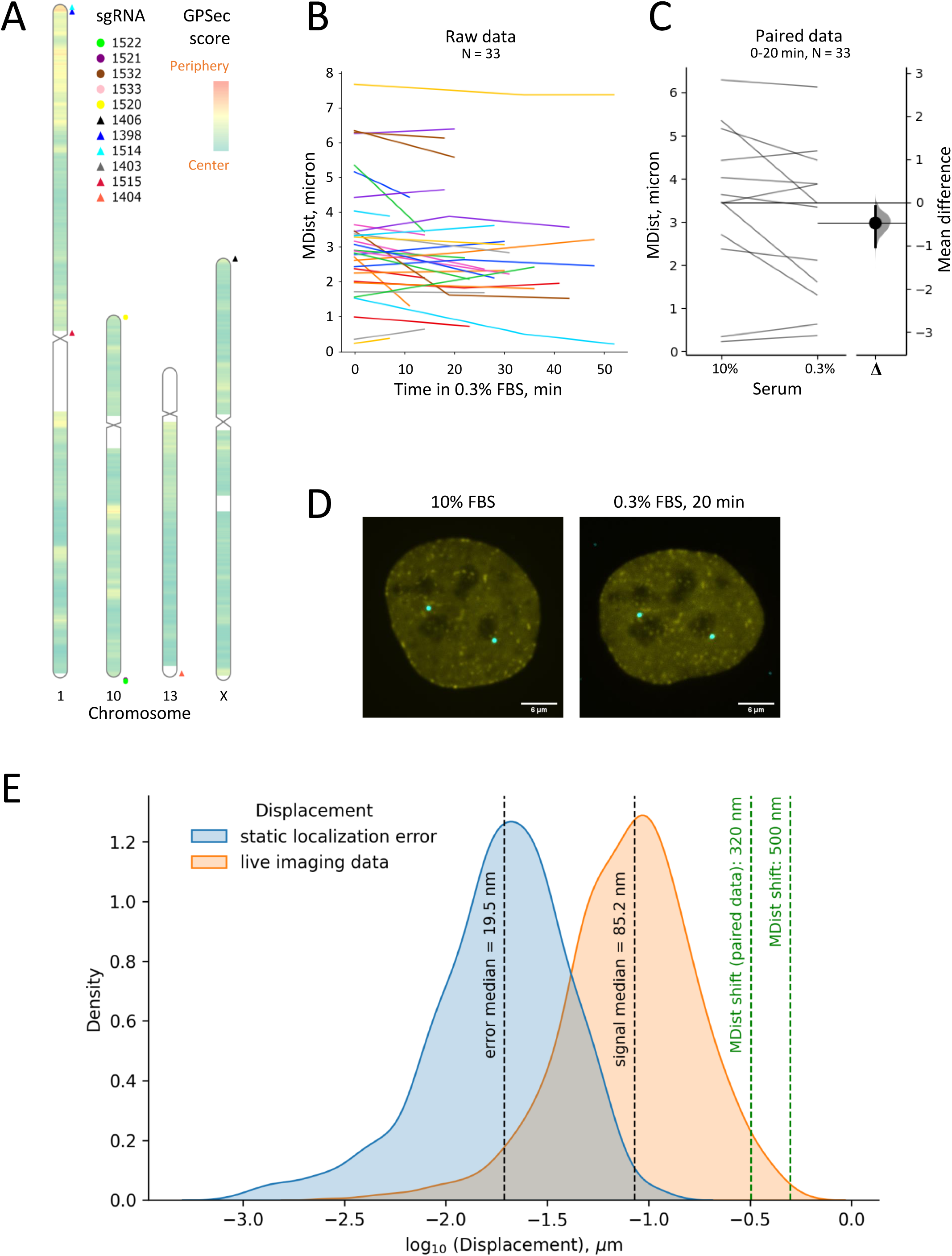
related to main figure 1. **A** Schematic of the chromosome localization of the target sequence sites labelled in this study with GPSec score (Girelli et al 2020) shown by color. Corresponding data with loci sequence and references can be found in Materials and Methods section. **B,C** Mean locus distance to nuclear periphery for chromosome 1 loci under starvation, measured in the same cell before and after medium change. **B** Raw data plotted without time-related subsampling, each line represents a nucleus. **C** Cells sampled within the first 20 min after medium change. Every line represents the change in minimal distance in particular locus, left scale. The distribution shows bootstrapped delta mean values for paired dataset, right scale. Error bars show 95% confidence intervals. **D** Representative image of U2OS cells overexpressing CRISPR- Sirius plasmids with guide for chromosome 1 locus, in 10% serum and under starvation. The descriptive statistics for cells imaged before and after the medium change is shown in Figure 1h. Scale bar = 6μm. **E.** Localization error determined as median of displacements of the set of immobile chromosome 1 loci in fixed cells (blue distribution) at imaging setup and processing pipeline identical to live experiment (orange distribution). Dashed lines show the change in minimal distance to nuclear edge observed in live cells under starvation, plotted on the same scale. Displacement length values are log10-transformed for clarity.

**Supplementary figure 4.**
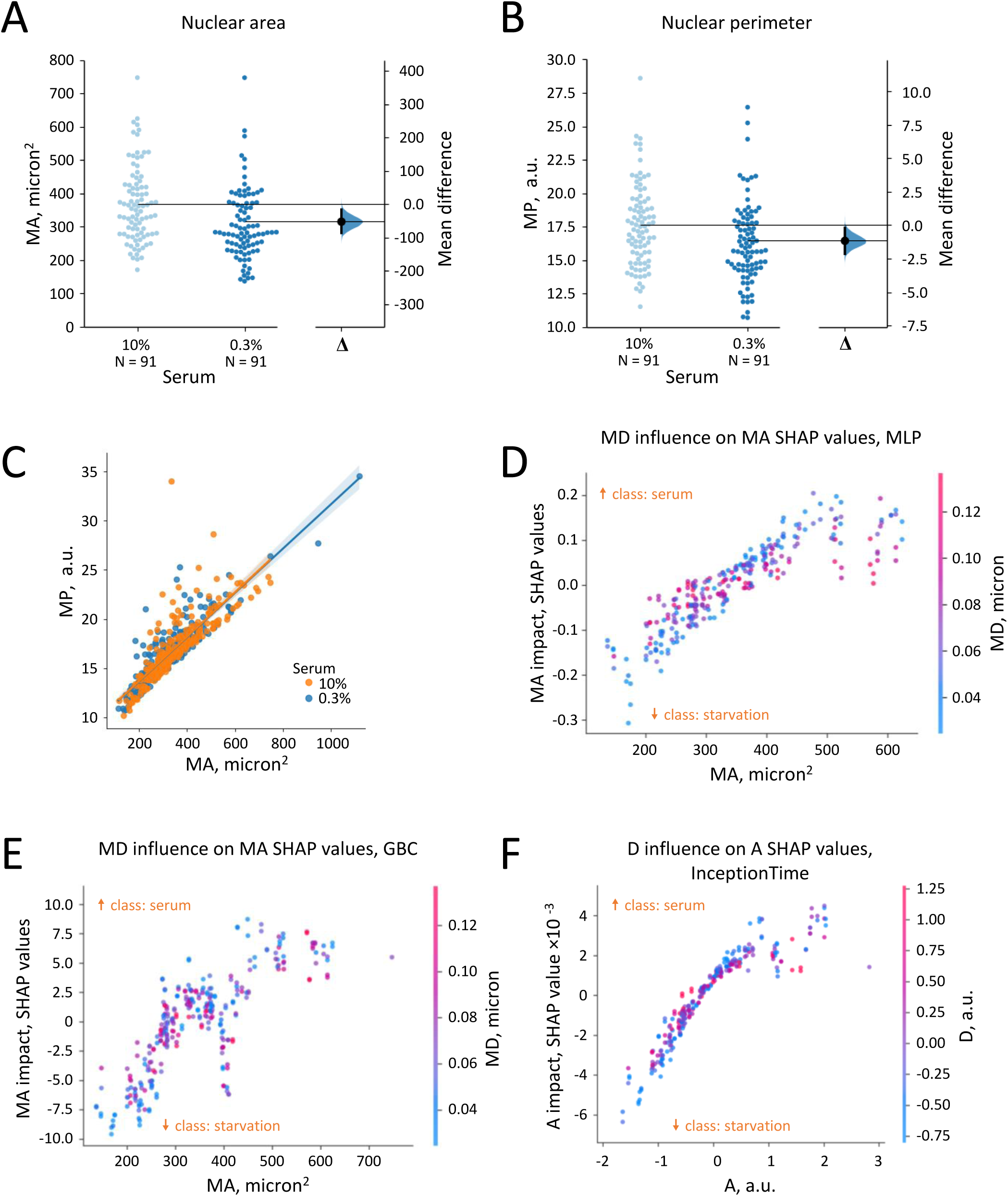
related to main figure 4. Changes in nuclear morphology under starvation, particularly, area (*MA*, **a**) and perimeter (*MP*, **b**). Each dot in the swarmplot represents an individual nucleus. Panels on the right represent the distributions of bootstrapped delta mean values for datasets. Error bars show 95% confidence intervals. **c**, Correlation between nuclear area (*MA*) and the length of perimeter (*MP*) in 10% FBS and in starvation (0.3% FBS). Shaded area shows 95% confidence interval. **d-f**, Locus speed as measured by mean locus displacement (*MD*) affects Shapley values for nuclear area (*MA*), which is clear from vertical color gradient on SHAP dependence plot, where the locus speed is color-mapped, see the legend on the right. SHAP was used to explain the decisions of MLP (**d**, repeated from Fig 4f for comparison), GBC (**e**), and InceptionTime (**f**) classifiers.

**Supplementary Table 1:** Inferential statistics results.

|  | Chromosome 1 |  | Telomeres |  |
| --- | --- | --- | --- | --- |
| Metric | MW | MW_pval | MW | MW_pval |
| MDist | **12946** | **0.04** | **27124** | **0.932** |
| mA | **14940** | **0** | n.a. | n.a. |
| sA | **13001** | **0.033** | n.a. | n.a. |
| MD | **10687** | **0.356** | 28312 | 0.365 |
| TD | **10781** | **0.424** | 23009 | 0.006 |
| DistR | **10290** | **0.148** | **21675** | **0** |
| TDist | 12378 | 0.192 | 26481 | 0.72 |

MW: Mann-Whitney U statistic, two-sided
MW\_pval: p-value
**Bold font** indicates that MW test results were in accordance with the descriptive statistics.

